# Puzzling parasitic plants: phylogenetics and classification of Santalales revisited

**DOI:** 10.1101/2025.05.16.654241

**Authors:** Luiz A. Cauz-Santos, James W. Byng, Mark W. Chase, Maarten J. M. Christenhusz

## Abstract

Based on a previously published but realigned matrix for Santalales, we find many relationships that were weakly or unsupported in previous studies are here much better supported, providing a more robust foundation upon which to discuss Santalales classification. In the maximum likelihood analysis, we recovered the same basic relationships as in the previous studies, but with two major differences: i) Balanophoraceae in the broad sense are monophyletic and well supported as embedded in Santalaceae (rather than biphyletic and outside Santalaceae) and ii) most of the former Olacaceae form a moderately supported clade (rather than a weakly supported grade). In the parsimony analysis, the position of Balanophoraceae s.l. is not well supported (although their broader circumscription is). We outline three possible options for a classification of the order and propose a new familial and subfamilial classification for Santalales. This hopefully provides a stable, user-friendly taxonomic framework that is phylogenetically well supported and more consistent with historical usage than some recently proposed systems, and provides taxa that can be more readily diagnosed morphologically. We recommend recognition of nine families (in phylogenetic sequence): Strombosiaceae, Erythropalaceae, Olacaceae, Opiliaceae, Santalaceae, Misodendraceae, Schoepfiaceae and Loranthaceae, plus an unresolved position for Balanophoraceae (including Mystropetalaceae), which we propose to exclude from Santalaceae until more evidence of their relationships to that family is available from nuclear genes. Four new subfamilies, Gaiadendroideae, Comandroideae, Nanodeoideae and Thesioideae, are proposed, and a new combination, *Loranthus obtusifolius*, is made.

## Introduction

In APG III and IV (2009, 2016), a few orders were highlighted as relatively problematic: e.g. Caryophyllales, Dioscoreales, Lamiales and most notably Santalales. No decision on a revised classification of the order Santalales was made, and further study was recommended (APG IV, 2016). These predominantly parasitic species have been difficult, because standard plastid markers do not always align well due to widespread insertions and deletions and highly divergent sequences. Sampling is also often an issue, and herbarium DNA is typically highly degraded in specimens that turn black when dried (as many of these do) or difficult to extract in a usable condition (e.g., Zuntini *et al*. 2024).

In APG IV (2016), Santalales were one of the last angiosperm orders with some clearly non-monophyletic families. The reasons behind the continued use of such familial concepts were that earlier analyses resorted to an approach that recommeneded too many small families without correlated morphological characters, making them difficult to identify (Nickrent *et al*. 2010, Su *et al*. 2015). To address this problem, we here re-evaluate the results of Su *et al*. (2015) with a new alignment. Based on these much better supported results, we outline three possible ways of classifying the order and propose one that recognises a smaller set of larger families, some of which admittedly still have no obvious morphological synapomorphies. However, our preference is for fewer, larger such morphologically “difficult” families.

Santalales comprises roughly 2522 species in 177 genera (Byng 2014, Kuijt & Hansen 2015, Christenhusz & Byng 2016, Christenhusz *et al*. 2017, POWO, 2025). In Santalales, Balanophoraceae comprise the only holoparasites. Most other genera are at least partially photosynthetic (i.e., hemiparasites), but several genera, e.g., *Erythropalum* Blume, *Scoridocarpus* Becc. and *Strombosia* Blume, in polyphyletic Olacaceae, are putatively autotrophic (Nickrent 1997, Nickrent *et al*. 2010, Byng 2014, Kuijt & Hansen 2015). Stem parasitism (the mistletoe habit, as opposed to root parasitism) has evolved independently several times, e.g., in Loranthaceae, Misodendraceae and Santalaceae.

Previous molecular phylogenetic studies have improved circumscription of Santalales, notably by excluding Dipentodontaceae (now Huerteales; Peng *et al*. 2003, Worburg et al. 2009, Christenhusz *et al*. 2010), Grubbiaceae (now Cornales; Xiang *et al*. 2002) and Medusandraceae (now Peridiscaceae, Saxifragales; Savolainen *et al*. 2000, Wurdack & Davis 2009). Like many holoparasitic plants, placement of Balanophoraceae has proven contentious due to its unusual morphology and substantially altered plastid genome (e. g., Bungard 2004). Phylogenetic studies by Nickrent *et al*. (2005), Barkman *et al*. (2007) and Su & Hu (2012) used 18S nuclear rDNA and mitochondrial *matR* and found Balanophoraceae to be associated with Santalales, but their position in the order was unclear. Some Santalales were included in the large angiosperm-wide, multi-nuclear gene analyses of Zuntini *et al*. (2024), but many critical taxa were not sampled, making this study inapplicable to the issue of familial classification in Santalaceae.

In APG IV (2016; unchanged from APG III 2009), seven families were recognised in Santalales: Balanophoraceae, Loranthaceae, Misodendraceae, Olacaceae, Opiliaceae, Santalaceae and Schoepfiaceae. However, it was noted then that Olacaceae and Santalaceae were not monophyletic (Der & Nickrent 2008, Malécot & Nickrent 2008, Malécot *et al*. 2004), a situation that requires a revised familial arrangement of some sort. APG IV (2016) rejected the tactic used in some published studies (see below), which involved recognising as families all well-supported clades despite not knowing their inter-relationships, an approach we describe here as the “least common denominator” strategy. If classification of families is to be altered, then their inter-relationships should not be highly speculative because other options, perhaps with more broadly defined families, cannot be considered.

Relationships in the order were assessed by various studies using different sets of species and markers (Nickrent & Malécot 2001, Vidal-Russell & Nickrent 2007, 2008a, Der & Nickrent 2008, Malécot & Nickrent 2008, Nickrent & Garcia 2015). However, relationships of many clades identified in these studies were unresolved, particularly in Olacaceae, which in the sense of APG III and IV (2009, 2016) were non-monophyletic (i.e., paraphyletic to the rest of the order, but with weak support; Nickrent & Malécot, 2001, Vidal-Russell & Nickrent 2008a, b). Despite poor resolution/support at many nodes, Nickrent *et al*. (2010) proposed a new familial classification for the order, naming all clades, resulting in recognition of 19 families. This lowest common denominator approach (Nickrent, Anderson & Kuijt 2019) meant that each well-supported clade was given a formal family name, despite lacking a clear picture of their inter-relationships.

Su *et al*. (2015) expanded the sampling of loci with a seven-gene analysis including three plastid genes (*matK, rbcL, accD*), one mitochondrial gene (*matR*) and three nuclear genes (SSU and LSU rDNA and *rpb2*) and increased sampling of genera of Santalales. They placed Balanophoraceae close to Loranthaceae. However, in this study Balanophoraceae were found to be biphyletic, splitting along subfamily lines, Balanophoroideae and Mystropetaloideae. Therefore, Su *et al*. (2015) proposed splitting them in two families: Balanophoraceae *s.s.* and Mystropetalaceae, which was supported by maximum likelihood and Bayesian analyses (Su *et al*. 2015: Fig. 1A-C: pp. 494–496), but weakly supported by parsimony (Su *et al*. 2015: Fig. S2).

**Figure 1.**
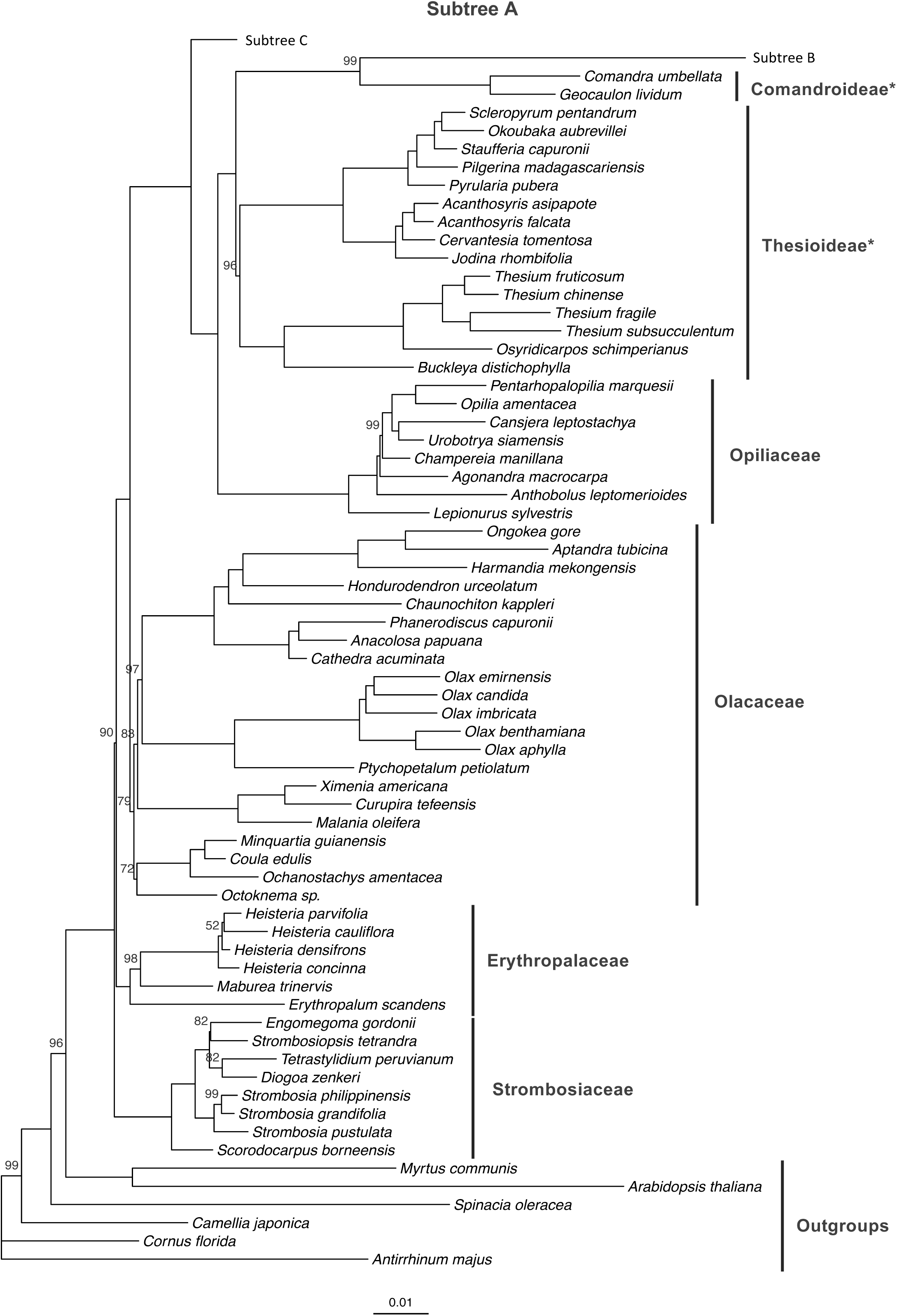
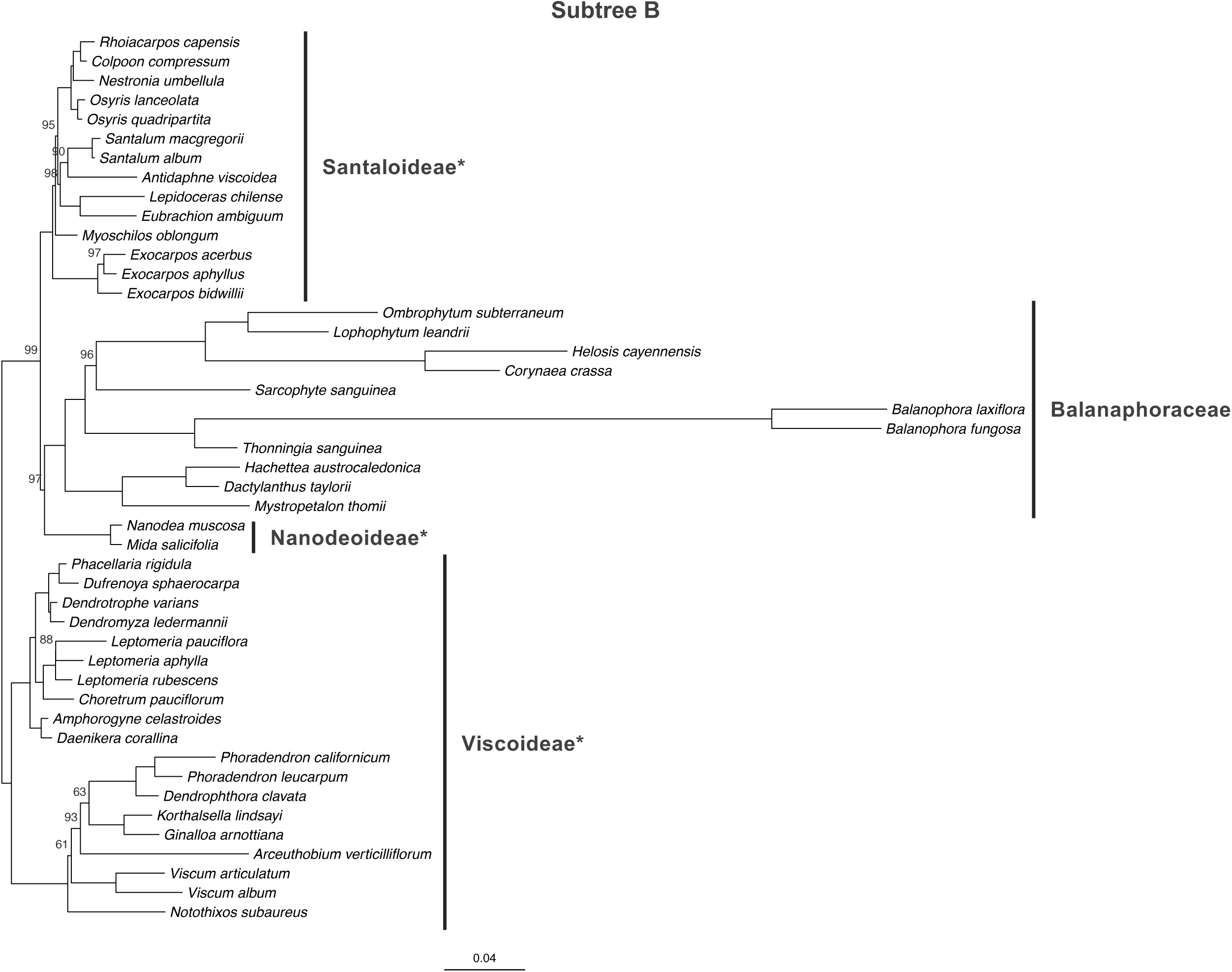
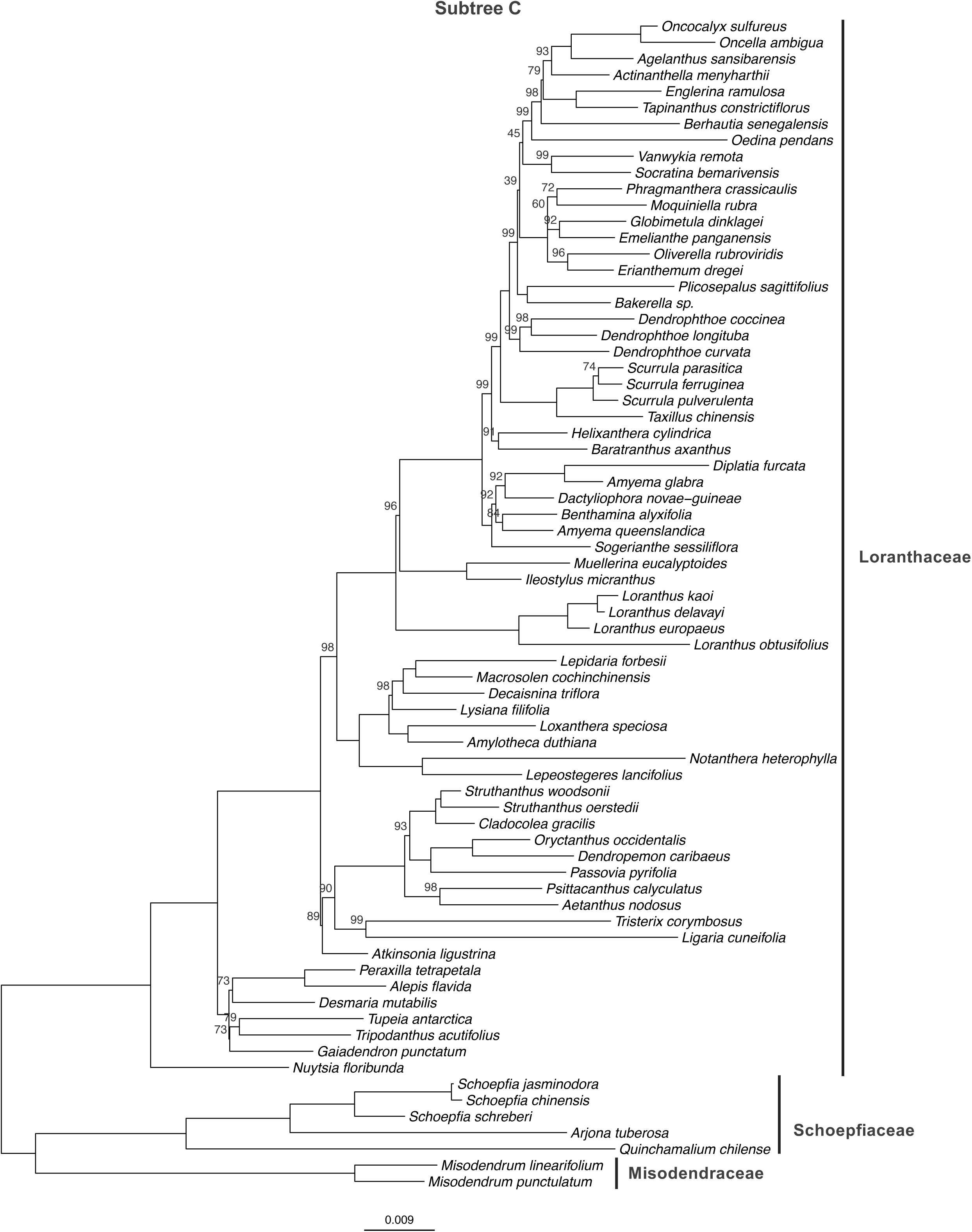
Maximum likelihood (ML) tree inferred from the seven-gene dataset for Santalales. ML bootstrap values are shown above branches; nodes with 100% support are not labeled. Our proposed classification is indicated on the right. Santalaceae subfamilies are marked with an asterisk (*).

Otherwise, the analyses of Su *et al*. (2015) corroborated the families of Nickrent *et al*. (2010), but as in the earlier analyses, interrelationships of these clades were less resolved and poorly supported. However, despite Amphorogynaceae, Viscaceae, Cervantesiaceae, Thesiaceae, Nanodeaceae, Santalaceae and Comandraceae forming a well-supported clade in the maximum likelihood and Bayesian trees, they did not consider expansion of Santalaceae (the oldest name) to include these, as was previously done.

The overall aim of this study is to provide an improved assessment of relationships in Santalales and re-examine relationships along the spine of the tree to provide a more robust classification supported by morphological synapomorphies. The Su *et al*. (2015) result perplexed us, and we decided to investigate their original matrix (provided by the authors as an electronic supplement). We found their matrix suffered from several alignment issues, which we have corrected and reanalysed below. Based on this improved phylogenetic analysis, we propose a more conservative classification with more, broader familial circumscriptions.

## Material & Methods

### Taxon sampling and gene selection

Sampling in this study (Table S1) included 184 accessions representing 149 genera (of 179, 83%) and all 19 previously recognised families *sensu* Nickrent *et al*. (2010) plus Mystropetalaceae *sensu* Su *et al*. (2015). Sequences for three nuclear (SSU and LSU ribosomal DNA and *rpb2*) and one mitochondrial (*matR*) and three plastid genes (*accD*, *matK*, *rbcL*) were downloaded from GenBank (Su *et al*. 2015).

### Phylogenetic analyses

All sequences for the seven-gene dataset were aligned individually using the multiple sequence alignment software MUSCLE (Edgar, 2004) implemented in *GeneiousPro* version 6.1.6 (http://www.geneious.com/; Biomatters Ltd., Auckland, New Zealand) using default settings. As for the Su *et al*. (2015) matrix, alignments were manually curated, for which we followed the guidelines of Kelchner (2000). We found that the aligment of the published Su *et al*. (2015) matrix (https://onlinelibrary.wiley.com/action/downloadSupplement?doi=10.12705%2F643.2&file=tax6432-sup-001-TXT.txt) introduced many unnecessary gaps, which decreased the number of phylogenetically informative positions. We combined the individual alignments in a single matrix and analysed this using maximum parsimony (MP) and maximum likelihood (ML) methods. The ML analysis was performed using IQ-TREE v.1.6.1 (Nguyen *et al*. 2015). The best-fit substitution model (GTR+F+I+R4) was selected automatically by ModelFinder (Kalyaanamoorthy *et al*. 2017) based on the Bayesian information criterion. Node support was estimated using 1,000 ultrafast bootstrap replicates (Hoang *et al*. 2018). The MP analysis was performed using PAUP* v.4.0b10 (Swofford 2002). Characters were unordered with equal weighting (Fitch 1971). A heuristic search was performed with random sequence addition for 1000 replicates using a tree bisection–reconnection (TBR) branch swapping algorithm, holding a maximum of ten trees per replicate. Node support was estimated with a bootstrap analysis using heuristic search and 1000 pseudoreplicates. Phylogenetic trees were visualized and edited in R using ape (Paradis *et al*. 2004), ggplot2 (Wickham 2016), ggtree (Yu *et al*. 2017), cowplot (Wilke 2024) and tidytree (Yu 2022). We did not conduct a Bayesian analysis as in Su *et al*. (2015) because their results were highly correlated with the ML analysis, both in terms of topology and support, thus making inclusion of ML and Bayesian trees mutually redundant.

To address the issue of different methods being used in our analyses versus those used by Su *et al*. (2015), we followed their protocol but used our realigned matrix. The ML analysis in Su *et al*. (2015) used Garli (genetic algorithm for rapid likelihood inference) v.2.0 (Zwickl, 2006) with 100 search replicates, stepwise addition of taxa and all other options on default settings. In RAxML v.7.0.4 (Stamatakis, 2006), Su *et al*. (2015) used 500 pseudo-replicates of rapid bootstrapping under the GTR + I + G model. Note also that we used 1000 pseudo-replicates of rapid bootstrapping under GTR+F+I+R4 model (as reported above).

## Results

### Matrix and result statistics

The final alignment comprised 12,637 characters (File S1) versus 12,884 in Su *et al*. (2015), of which 5,039 were potentially parsimony-informative versus 4,674 in Su *et al*. (2015), an increase of 9.4%. The ML analysis produced a well-resolved tree with high support across most nodes. The best ML tree had a log-likelihood of -162,143.57 (s.e. 1,822.83) and a total tree length of 46,154.

This tree had 182 nodes, with 123 (67.6%) receiving 100 bootstrap percentage (BP) and 38 (20.9%) between BP 90–99. In the MP analysis, the tree length was 27,206 steps with a consistency index (CI) of 0.41 and a retention index (RI) of 0.74, both better than those reported in Su *et al*. (2015; CI 0.38 and RI 0.64).

### Phylogenetic results

Although generally consistent with the Su *et al*. (2015) results, our combined analysis of the same seven-gene dataset recovered slightly different results with improved resolution and bootstrap support (Fig. 1). Overall, 167 nodes (91.8%) in our ML tree had 80% or higher, whereas in the ML results in Su *et al*. (2015) only ca. 70% of the nodes had ML bootstrap support higher than 80%. Our MP tree (Fig. S1) is in general like the ML tree, but with lower support and resolution. Similar parsimony results were also noted by Su *et al*. (2015). Below we report solely the ML topology and bootstrap percentages. Our favoured option for the classification of Santalales paralleles that of Nickrent *et al*. (2010), but we use below and in the illustrated tree our proposed subfamily names corresponding to some of their families to facilitate comparsison to the results in Su *et al*. (2015).

Olacaceae *sensu* APG III & VI (2009, 2016) form a grade in the ML tree with the other families embedded (Fig. 1). This consists of unresolved Strombosiaceae and Erythropalaceae (both well supported, BP 100) as sister (BP 100) to all other members of the order. A clade representing the rest of Olacaceae (including Aptandraceae, Coulaceae, Octoknemaceae, Olacaceae and Ximeniaceae sensu Nickrent et al. (2010) (each individually BP 100 except for Octoknemaceae, which has only a single accession) was moderately supported (BP 83). This clade was sister to the rest of Santalales (BP 100), which comprises all ‘non-olacoid’ taxa comprising two subclades. In one are Loranthaceae (BP 100; see below) as sister (BP 100) to Misodendraceae (BP 100) plus Schoepfiaceae (BP 100). In the other subclade, Opiliaceae (BP 100) are well-supported as sister (BP 100) to the rest of the families, including Balanophoraceae (plus Mystropetalaceae BP 100) sister (BP 97) to Nanodeoideae (BP 100; Fig. 1). Balanophoraceae plus Nanodeoideae are sister (BP 99) to Santaloideae (BP 100). This clade (BP 99) is then sister (BP 100) to Comandroideae (BP 100). Finally, Viscoideae (BP 100; including Amphorogynaceae sensu Nickrent *et al*. 2010) are sister (BP 100) to all but Opiliaceae. We note that although we found Balanophoraceae s.l. to be weakly supported as monophyletic in the parsimony tree (BP 71), they are only weakly supported (BP 68) as sister to Nanodeoideae, Santaloideae and Viscoideae.

We will not provide commentary on most relationships in Loranthaceae because this is outside the focus of this paper, but we note that *Amyema* Tiegh. and *Helixanthera* Lour. appear to be polyphyletic. The polyphyly of the latter can be resolved by placing *Helixanthera coccinea* (Jack) Danser in *Dendrophthoe* Mart. (as *Dendrophthoe coccinea* (Jack) G.Don). *Cecarria* Barlow is shown to be closely related to *Loranthus* Jacq. and may be included in that genus on morphological and phylogenetic grounds [***Loranthus obtusifolius*** (Merr.) Cauz, Byng, M.W.Chase & Christenh., **comb. nov.** Basionym: *Phrygilanthus obtusifolius* Merr. in Philipp. J. Sci. 1(Suppl.): 189 (1906)]. Further study of generic circumscription in Loranthaceae is critically needed.

In the ML analysis using Garli with a different model and fewer pseudorelicates in the bootstrap analyses than in Su *et al*. (2015), we obtained results almost identical to those described above (i.e., more like our initial results than to those in Su *et al*. 2015), but with lower bootstrap support at some nodes (Fig. S2).

The most important difference is that the node supporting a portion of Olacaceae as monophyletic (see below) received BP <50, whereas with our methods and a realigned matrix this node received BP 83. Balanophoraceae s.l. are still monophyletic and highly supported (BP 100) as embedded in Santalaceae (BP 98).

## Discussion

Phylogenetic relationships among the major clades in Santalales found here are generally congruent with those presented in Su *et al*. (2015), with the notable exception of Balanophoraceae and Mystropetalaceae, which here in the ML tree are well supported as sister to Nanodeoideae (Fig. 1). Su *et al*. (2015) explained the diphyly of Balanophoraceae by stating that they differ widely in substitution rates, Balanophoraceae *s.s.* rapidly evolving versus Mystropetalaceae relatively more slowly. They also found that Balanophoraceae were monophyletic in their MP tree but with low support, whereas in their ML analysis Mystropetalaceae were sister to Loranthaceae with moderate support. This result is almost certainly related to alignment issues; we checked their alignment, which introduced gaps that removed nearly 10% of the potentially informative positions (see new aligment in our Suppl. File S1). Our alignment produced trees with a higher CI and RI, which indicated that our reduction in the number of introduced gaps did not increase “noise” in the analysis. A poor alignment has been documented to eliminate evidence of relationships (Philippe *et al*. 2011). The increased support along the spine of the tree in our ML tree underscores the importance of alignment and demonstrates that introducing too many gaps can eliminate what is otherwise clear evidence of relationships. We also used a different model from that in Su *et al*. (2015) and more bootstrap pseudoreplicates, which appears to have reduced bootstrap support for some nodes in the latter, but the topology is almost identical, leaving us the impression that the major cause of the topological differences was the introduction of too many unnecessary gaps. Some differences in support may also perhaps be due to a different model and fewer bootstrap replicates in Su et al. (2015).

The classification of Nickrent *et al*. (2010) was based on several independent studies focussing on traditionally recognised families, such as Olacaceae *s.l*. (Malécot & Nickrent, 2008, Malécot et al., 2004), Loranthaceae (Vidal-Russell & Nickrent, 2008b) and Santalaceae *s.l.* (Der & Nickrent, 2008). Until Su *et al*. (2015), no comprehensive analysis for the order had been published, and thus the classification of Nickrent *et al*. (2010) was premature. Once their classification was published, Nickrent and colleagues persisted in recognising smaller, difficult to circumscribe families, despite evidence in Su *et al*. (2015) of the presence of some well-supported larger groupings more consistent with previously recognised families. In particular, Su *et al*. (2015) found a well-supported clade comprising Amphorogynaceae, Cervantesiaceae, Comandraceae, Nanodeaceae, Santalaceae and Viscaceae, but they did not consider enlarging Santalaceae to include these other families.

The more recent classification in Kuijt & Hansen (2015) ignored the molecular results and established known relationships. For example, they placed *Anthobolus* R.Br. and *Arjona* Comm. ex Cav. + *Quinchamalium* Molina in Santalaceae s.l., rather than in Opiliaceae and Schoepfiaceae, respectively (POWO, 2025), where they already had been placed based on molecular data. The phylogenetic relationships of Su *et al*. (2015) do correspond well with the classification proposed by Nickrent *et al*. (2010). Despite some improvements in support and resolution, there was weak support for deeper nodes in the order, particularly the clades in the paraphyletic grade formed by the Olacaceae *s.l.* (e.g. *Aptandra*, *Coula*, *Erythropalum*, *Octoknema*, *Olax*, *Strombosia* and *Ximenia* groups) and placement of holoparasitic Balanophoraceae.

This was the reasoning why no changes were made to the APG III (2009) classification of this order when proposing an updated classification in APG IV (2016).

In the nuclear phylogenetic tree in Zuntini *et al*. (2024), a different topology is found, but most of this probably results from issues with the holoparasitic Balanophoraceae, which they found widely polyphyletic. The backbone of Santalales in that study also exhibits low support, and Olacaceae suffers from a paucity of sampling, but similarly formed a grade to the remaining Santalaceae.

Our re-analysis of the Su *et al*. (2015) matrix found good support for the *Strombosia* clade (Strombosiaceae sensu Nickrent *al.* 2010) and the *Erythropalum* clade (Erythropalaceae sensu Nickrent *al.* 2010), although it remains unclear how these relate to each other. The deeper relationships among the remaining olacoid clades are only moderately supported in our analysis, but this is an improvement over the results in Su *et al*. (2015) in which these relationships were unresolved. Unlike Su *et al*. (2015), Balanophoraceae and Mystropetalaceae here form a well-supported clade deeply embedded in the Santalaceae clade in the ML tree.

### Classification

Nickrent *et al*. (2010) more than doubled the number of families in the order, and as the authors themselves stated in their revised classification the recognised families “are, in some cases, more difficult to identify and circumscribe using morphological features”. For example, separating Santalaceae *s.s.* from Amphorogynaceae is impossible on morphological grounds and can only be done by identifying the genus and then looking up to which family that genus now belongs (Byng 2014). The Santalales treatment of Kuijt & Hansen (2015) diverged substantially from that of Nickrent *et al*. (2010), but molecular results were not considered in this study and thus several genera were placed in families where they do not belong. Molecular studies have shown that previous morphological classifications often do not reflect evolutionary relationships in Santalales (Nickrent & Malécot, 2001; Vidal-Russell & Nickrent, 2007, 2008a; Der & Nickrent, 2008; Sun *et al*. 2015

APG IV (2016) opted for making no changes to the family classification of Santalales and followed APG III (2009), even though it was known to the APG compilers at the time that Olacaceae and Santalaceae in this circumscription were not monophyletic. They stated that further molecular work should be undertaken to form the basis of a new classification with the hope that adding more data and taxa would clarify inter-relationships of the Nickrent *et al*. (2010) families, making other options possible (e.g., broader families more consistent with historical usage).

Taking our results into consideration, three options are considered here (Fig. 1; Table 1):

1. Accept all families proposed by Nickrent *et al*. (2010), increasing the number of families in the order to 19 (if Balanophoraceae s.l. are accepted). This includes around ten families that are mostly morphologically difficult or impossible to diagnose and mostly with few (1–5) genera. This option leaves open the possibility of additional families being described if future analysis with more complete taxonomic sampling demonstrates that some of these 19 families are not monophyletic. This option also deviates substantially from all previous taxonomic treatments of Santalales, although to follow this option, no new families will need to be described (they were described in Nickrent *et al*. 2010).
2. Based on clearer relationships, recognition of fewer but largely morphologically circumscribable broader families. This will nonetheless involve a few families that are morphologically similar, and others that are morphologically diverse. It maintains many clade concepts of Nickrent *et al*. (2010) but relegates them to subfamily status. This would include the seven families of APG IV (Balanophoraceae, Loranthaceae, Misodendraceae, Olacaceae, Opiliaceae, Santalaceae s.l. and Schoepfiaceae) plus Erythropalaceae and Strombosiaceae. Balanophoraceae can either be treated as part of Santalaceae (as a separate subfamily) based on the well supported embedded position found here, or maintained apart from Santalaceae s.l. (including Cervantesiaceae Comandraceae, Nanodeaceae, Thesiaceae and Viscaceae, the last including Amphorogynaceae) due to the known liabilities of placing parasitic taxa in phylogenetic studies involving plastid genes, which provide a large portion of the data used in this study. In the future, when more nuclear data become available, it is possible that some reorganisation in Santalaceae s.l. may be needed, especially concerning the position of Balanophoraceae, but this will not require major name changes across the board and thus this option is likely to be most stable in the long run.
3. Accept only a single family in the entire order but providing a series of subfamilies that will be easily recognisable morphologically (e.g., mainly parasitic, sepals often reduced, petals often valvate, fruits usually indehiscent, leaf margins entire, stipules absent, carpels usually three and fused, albeit in diverse ways). This option would be extremely stable as no other family classification would be needed, and future changes can be dealt with at the subfamilial level. This would be the most stable, long-term option, but it will be destabilising initially because Loranthaceae with such broad circumscription has no precedent.

**Table 1.**
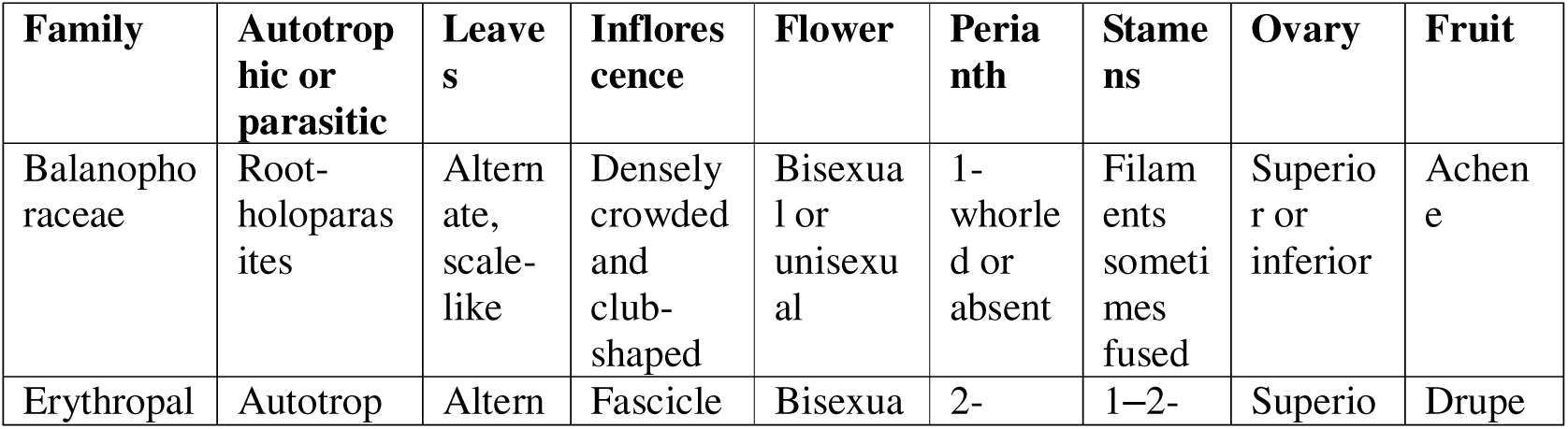
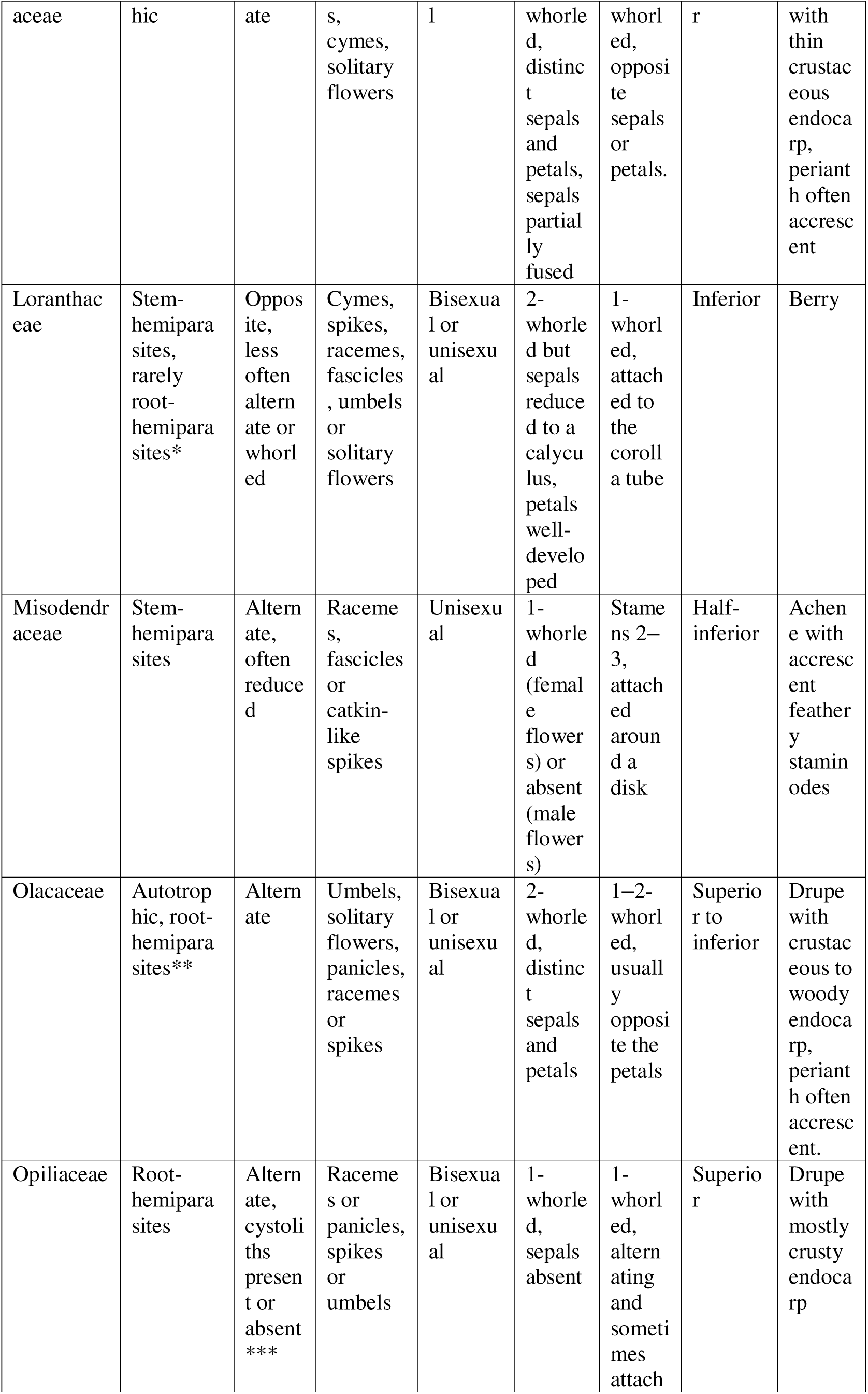
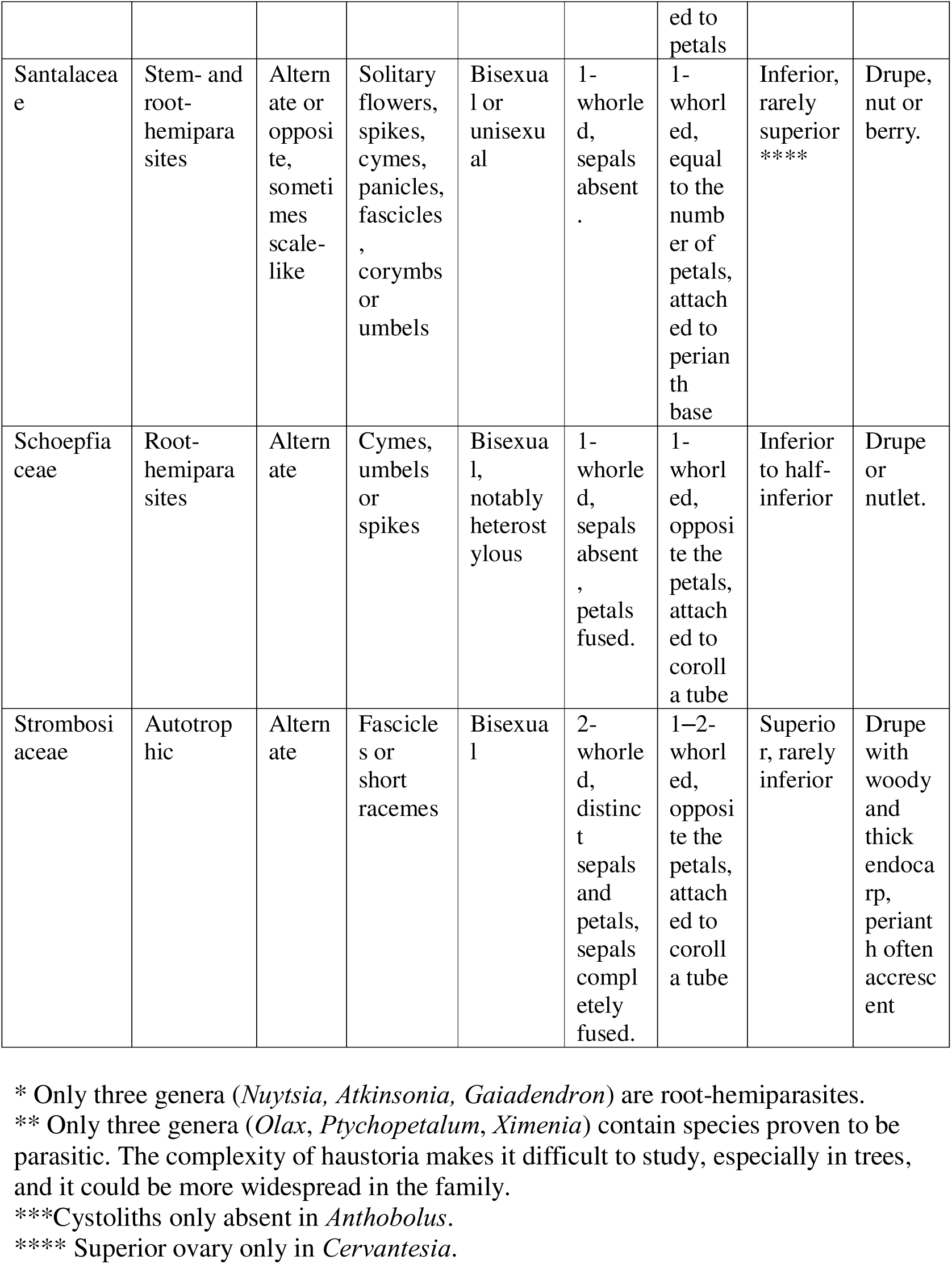
Macromorphological characters compared of Santalales families. Some key characters have some notable exceptions listed.

We propose option 2 for the classification of Santalales and provide a list of synapomorphies for the families (Table 1). This is the option most consistent with the APG III and APG IV treatments, and, given the broader family limits, it should also be relatively robust to changes necessitated by future studies with better generic representation.

### Proposed classification of Santalales

(*genus placement based on unpublished analyses; † genus not sampled; species numbers from POWO, 2025).

**Balanophoraceae** Engl., *incertae sedis* in Santalales (16 genera, 60 species)

***Balanophora*** J.R.Forst. & G.Forst. (25 spp.)
***Chlamydophytum*** Mildbr. (1 sp.)†
***Corynaea*** Hook.f. (1 sp.)
***Dactylanthus*** Hook.f. (1 sp.)
***Ditepalanthus*** Fagerl. (2 sp.)†
***Hachettea*** Baill. (1 sp.)
***Helosis*** Rich. (3 spp.)
***Langsdorffia*** Mart. (3 spp.)†
***Lathrophytum*** Eichler (1 sp.)†
***Lophophytum*** Schott & Endl. (4 spp.)
***Mystropetalon*** Harv. (1 sp.)
***Ombrophytum*** Poepp. ex Endl. (7 spp.)
***Rhopalocnemis*** Jungh. (1 sp.)†
***Sarcophyte*** Sparrm. (2 spp.)
***Scybalium*** Schott & Endl. (4 spp.)†
***Thonningia*** Vahl (3 spp.)

1) **Strombosiaceae** Tiegh. (6 genera, 20 species)

***Diogoa*** Exell & Mendonça (2 spp.)
***Engomegoma*** Breteler (1 sp.)
***Scorodocarpus*** Becc. (1 sp.)
***Strombosia*** Blume (11 spp.)
***Strombosiopsis*** Engl. (3 spp.)
***Tetrastylidium*** Engl. (2 spp.)

2) **Erythropalaceae** Planch. ex Miq., nom. cons. (4 genera, 43 species)

***Brachynema*** Benth. (2 spp.)*
***Erythropalum*** Blume (1 sp.)
***Heisteria*** Jacq. (39 spp.)
***Maburea*** Maas (1 sp.)

3) **Olacaceae** R.Br., nom. cons. (incl. Aptandraceae, Coulaceae, Octoknemaceae, Ximeniaceae) (18 genera, 121 species). Note: *Octoknema* is problematic due to the great number of autapomorphies exhibited: “They are members of Olacaceae (s.l.), but are unusual in having a stellate indumentum, being dioecious, and having endosperm lamina intrusive into the deeply ruminate seeds”; Gosline & Malecot 2011).

***Anacolosa*** (Blume) Blume (16 spp.)
***Aptandra*** Miers (4 spp.)
***Cathedra*** Miers (6 spp.)
***Chaunochiton*** Benth. (3 spp.)
***Coula*** Baill. (1 sp.)
***Curupira*** G.A.Black (1 sp.)
***Douradoa*** Sleumer (1 sp.)†
***Harmandia*** Pierre ex Baill. (1 sp.)
***Hondurodendron*** C.Ulloa, Nickrent, Whitef. & D.L.Kelly (1 sp.)
***Malania*** Chun & S.K.Lee (1 sp.)
***Minquartia*** Aubl. (1 sp.)
***Ochanostachys*** Mast. (1 sp.)
***Octoknema*** Pierre (14 spp.)
***Olax*** L. (incl. *Dulacia* Vell.) (52 spp.)
***Ongokea*** Pierre (1 sp.)
***Phanerodiscus*** Cavaco (3 spp.)
***Ptychopetalum*** Benth. (4 spp.)
***Ximenia*** Plum. ex L. (10 spp.)

4) **Opiliaceae** Valeton, nom. cons. (11 genera, 38 species)

***Agonandra*** Miers ex Benth. (10 spp.)
***Anthobolus*** R.Br. (4 spp.)
***Cansjera*** Juss. (3 spp.)
***Champereia*** Griff. (1 sp.)
***Gjellerupia*** Lauterb. (1 sp.)†
***Lepionurus*** Blume (1 sp.)
***Melientha*** Pierre (1 sp.)†
***Opilia*** Roxb. (2 spp.)
***Pentarhopalopilia*** (Engl.) Hiepko (4 spp.)
***Rhopalopilia*** Pierre (3 spp.)†
***Urobotrya*** Stapf (8 spp.)

5) **Santalaceae** R.Br., nom. cons. (incl. Amphorogyne, ’’, ’Cervantesiaceae’, ’Comandraceae’, ’Mystropetalaceae’, ’Nanodeaceae’, ’Thesiaceae’, ’Viscaceae’) (57 genera, 1192 species)

**Subfamily Thesioideae** (Vest) Cauz, Byng, M.W.Chase & Christenh., **stat. nov**. Basionym: Thesiaceae Vest, Anleit. Stud. Bot.: 270, 289. 1818 (*Thesioideae*).

Validated by a description in German. – T: *Thesium* L. (1753). (Thesiaceae + Cervantesiaceae)

***Acanthosyris*** (Eichler) Griseb. (6 spp.)
***Buckleya*** Torr. (5 spp.)
***Cervantesia*** Ruiz & Pav. (2 spp.)
***Jodina*** Hook. & Arn. ex Meisn. (1 sp.)
***Okoubaka*** Pellegr. & Normand (2 spp.)
***Osyridicarpos*** A.DC. (1 sp.)
***Pilgerina*** Z.S.Rogers, Nickrent & Malécot (1 sp.)
***Pyrularia*** Michx. (2 spp.)
***Scleropyrum*** Arn. (5 spp.)
***Staufferia*** Z.S.Rogers, Nickrent & Malécot (1 sp.)
***Thesium*** L. (incl. *Kunkeliella* Stearn) (345 spp.)

**Subfamily Comandroideae** (Nickrent & Der) Cauz, Byng, M.W.Chase & Christenh., **stat. nov**. Basionym: Comandraceae Nickrent & Der in D.L. Nickrent et al., Taxon 59: 550. 4 Apr 2010. Validated by a description in Latin. – Type: *Comandra* Nutt. (1818).

***Comandra*** Nutt. (1 sp.)

***Geocaulon*** Fernald (1 sp.)

**Subfamily Viscoideae** Eichler (’Amphorogynaceae’ + ’Viscaceae’)
***Amphorogyne*** Stauffer & Hürl. (3 spp.)
***Arceuthobium*** M.Bieb. (38 spp.)
***Choretrum*** R.Br. (6 spp.)
***Daenikera*** Hürl. & Stauffer (1 sp.)
***Dendromyza*** Danser (30 spp.)
***Dendrophthora*** Eichler (138 spp.)
***Dendrotrophe*** Miq. (incl. *Henslowia* Blume) (5 spp.)
***Dufrenoya*** Chatin (incl. *Hylomyza* Danser) (13 spp.)
***Ginalloa*** Korth. (9 spp.)
***Korthalsella*** Tiegh. (29 spp.)
***Leptomeria*** R.Br. (incl. *Spirogardnera* Stauffer) (18 spp.)
***Notothixos*** Oliv. (8 spp.)
***Phacellaria*** Benth. (6 spp.)
***Phoradendron*** Nutt. (271 spp.)
***Viscum*** L. (112 spp.)

**Subfamily Nanodeoideae** (Nickrent & Der) Cauz, Byng, M.W.Chase & Christenh., **stat. nov**. Basionym: Nanodeaceae Nickrent & Der in D.L. Nickrent et al., Taxon 59: 1. 552. 4 Apr 2010. Validated by a description in Latin. – T: *Nanodea* Banks ex C.F. Gaertn. (1807).

**Mida** R.Cunn. ex A.Cunn. (1 sp.)

**Nanodea** Banks ex C.F.Gaertn. (1 sp.)

**Subfamily Santaloideae** Arn.

***Antidaphne*** Poepp. & Endl. (9 spp.)
***Colpoon*** P.J.Bergius (2 spp.)
***Eubrachion*** Hook.f. (2 spp.)
***Exocarpos*** Labill. (incl. *Elaphanthera* N.Hallé, *Omphacomeria* A.DC. (29 spp.)
***Lacomucinaea*** Nickrent & M.A.García (1 sp.)†
***Lepidoceras*** Hook.f. (2 spp.)
***Myoschilos*** Ruiz & Pav. (1 sp.)
***Nestronia*** Raf. (1 sp.)
***Osyris*** L. (3 spp.)
***Rhoiacarpos*** A.DC. (1 sp.)
***Santalum*** L. (19 spp.)

6) **Misodendraceae** J.Agadh., nom. cons. (1 genus, 8 species)

***Misodendrum*** Banks ex DC. (8 spp.)

7) **Schoepfiaceae** Blume (3 genera, 34 species)

***Arjona*** Cav. (5 spp.)
***Quinchamalium*** Molina (1 sp.)
***Schoepfia*** Schreb. (28 spp.)

8) **Loranthaceae** Juss., nom. cons. (77 genera, 1075 species)

Subfamily Nuytsioideae Tiegh.

***Nuytsia*** R.Br. ex G.Don (1 sp.)

**Subfamily Gaiadendroideae** (Tiegh. ex Nakai) Cauz, Byng, M.W.Chase & Christenh., **stat. nov**. Basionym: Gaiadendraceae Tiegh. ex Nakai in Bull. Natl. Sci. Mus. Tokyo 31: 45. Mar 1952 (Giadendraceae). Validated by a diagnosis in Latin. – T: Gaiadendron G. Don (1834).

***Alepis*** Tiegh. (1 sp.)
***Desmaria*** Tiegh. (1 sp.)
***Gaiadendron*** G.Don (2 spp.)
***Peraxilla*** Tiegh. (3 spp.)
***Tripodanthus*** Tiegh. (3 spp.)
***Tupeia*** Cham. & Schltdl. (1 sp.)

**Subfamily Loranthoideae** Eaton

***Actinanthella*** Balle (2 spp.)
***Aetanthus*** (Eichler) Engl. (17 spp.)
***Agelanthus*** Tiegh. (59 spp.)
***Amyema*** Tiegh. (94 spp.), polyphyletic
***Amylotheca*** Tiegh. (5 spp.)
***Atkinsonia*** F.Muell. (1 sp.)
***Bakerella*** Tiegh. (16 spp.)
***Barathranthus*** (Korth.) Miq. (4 spp.)
***Benthamina*** Tiegh. (1 sp.), embedded in *Amyema*
***Berhautia*** Balle (1 sp.)
***Cladocolea*** Tiegh. (29 spp.)
***Cyne*** Danser (7 spp.)†
***Dactyliophora*** Tiegh. (2 spp.), embedded in *Amyema*
***Decaisnina*** Tiegh. (27 spp.)
***Dendropemon*** (Blume) Rchb. (33 spp.)
***Dendrophthoe*** Mart. (incl. *Helixanthera coccinea* (Jack) Danser) (41 spp.)
***Diplatia*** Tiegh. (3 spp.), embedded in *Amyema*
***Distrianthes*** Danser (2 spp.)†
***Elytranthe*** (Blume) Blume (4 spp.)†
***Emelianthe*** Danser (1 sp.)
***Englerina*** Tiegh. (26 spp.)
***Erianthemum*** Tiegh. (17 spp.)
***Globimetula*** Tiegh. (13 spp.)
***Helicanthes*** Danser (1 sp.)
***Helixanthera*** Lour. (43 spp.)
***Ileostylus*** Tiegh. (1 sp.)
***Lampas*** Danser (1 sp.)
***Lepeostegeres*** Blume (10 spp.)
***Lepidaria*** Tiegh. (9 spp.)
***Ligaria*** Tiegh. (2 spp.)
***Loranthella*** S.Blanco & C.E.Wetzel (1 sp.)
***Loranthus*** Jacq. (incl. *Cecarria* Barlow) (8 spp.)
***Loxanthera*** (Blume) Blume (1 sp.)
***Lysiana*** Tiegh. (8 spp.)
***Macrosolen*** (Blume) Rchb. (43 spp.)
***Maracanthus*** Kuijt (3 spp.)†
***Moquiniella*** Balle (1 sp.)
***Muellerina*** Tiegh. (5 spp.)
***Notanthera*** G.Don (1 sp.)
***Oedina*** Tiegh. (4 spp.)
***Oliverella*** Tiegh. (3 spp.)
***Oncella*** Tiegh. (4 spp.)
***Oncocalyx*** Tiegh. (12 spp.)
***Oryctanthus*** Eichler (18 spp.)
***Oryctina*** Tiegh. (4 spp.)†
***Panamanthus*** Kuijt (1 sp.)†
***Papuanthes*** Danser (1 sp.)†
***Passovia*** H.Karst. (24 spp.)†
***Pedistylis*** Wiens (1 sp.)†
***Peristethium*** Tiegh. (18 spp.)†
***Phragmanthera*** Tiegh. (36 spp.)
***Phthirusa*** Mart. (18 spp.)
***Plicosepalus*** Tiegh. (12 spp.)
***Psittacanthus*** Mart. (131 spp.)
***Pusillanthus*** Kuijt (2 spp.)
***Scurrula*** L. (27 spp.)
***Septemeranthus*** L.J.Singh (1 sp.)†
***Septulina*** Tiegh. (2 spp.)†
***Socratina*** Balle (3 spp.)
***Sogerianthe*** Danser (6 spp.)
***Spragueanella*** Balle (2 spp.)
***Struthanthus*** Mart. (96 spp.)
***Tapinanthus*** (Blume) Rchb. (30 spp.)
***Taxillus*** Tiegh. (35 spp.)
***Thaumasianthes*** Danser (1 sp.)†
***Tolypanthus*** (Blume) Rchb. (7 spp.)†
***Trilepidea*** Tiegh. (1 sp.)†
***Tristerix*** Mart. (13 spp.)
***Trithecanthera*** Tiegh. (5 spp.)†
***Vanwykia*** Wiens (2 spp.)

## Acknowledgenents

Luiz A. Cauz-Santos was supported by a Marie Skłodowska-Curie Individual Fellowship under the CondensDrought project (Grant Agreement No. 101029312).

## Supplementary Materials

**Figure S1.** Maximum parsimony (MP) tree inferred from the seven-gene dataset for *Santalales*. Bootstrap values are shown above branches; nodes with 100% support are not labeled. Our proposed classification is indicated on the right. Subfamilies of *Santalaceae* are marked with an asterisk (*).

**Figure S2.** Maximum likelihood (ML) tree inferred from the seven-gene dataset for *Santalales*. To address differences in phylogenetic inference methods between our study and Su et al. (2015), we followed their ML analysis protocol (using Garli v.2.0 and RAxML v.7.0.4) but applied it to our realigned matrix. Bootstrap values are shown above branches; nodes with 100% support are not labeled. Our proposed classification is indicated on the right. Subfamilies of *Santalaceae* are marked with an asterisk (*).

**Table S1.** Taxa used in the phylogenetic analyses. GenBank accession numbers are provided for the following markers in this order: *SSU rDNA*, *LSU rDNA*, *RPB2*, *rbcL*, *matK*, *accD*, and *matR*. A dash (–) indicates that the corresponding sequence is missing. Table adapted from Su et al. (2015).

**File S1.** Alignment matrix used in the phylogenetic analyses, based on the seven-gene dataset for *Santalales*.

