## Supplementary figures and images for "Puzzling parasitic plants: phylogenetics and classification of Santalales revisited"

### Fig. S1

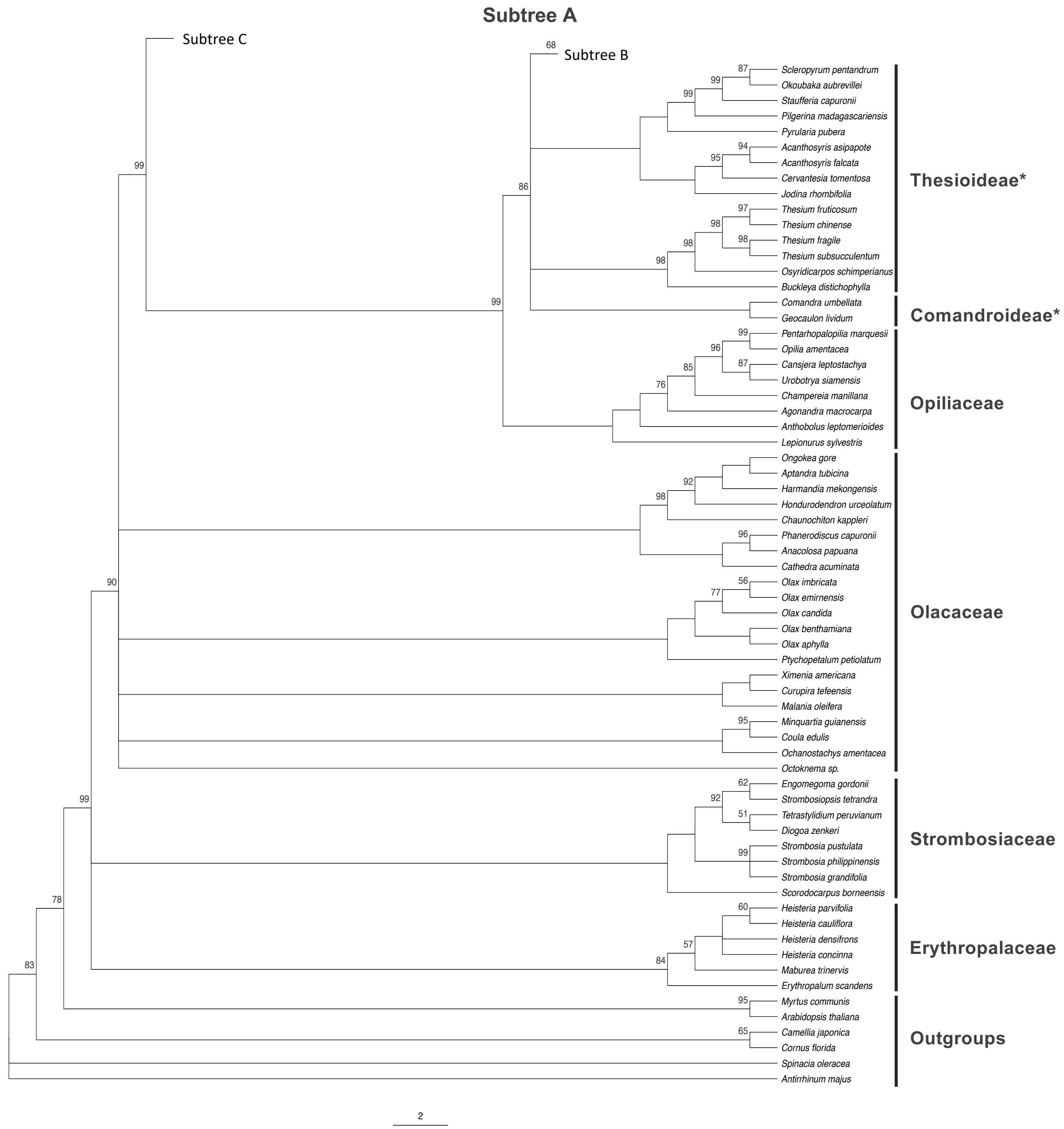

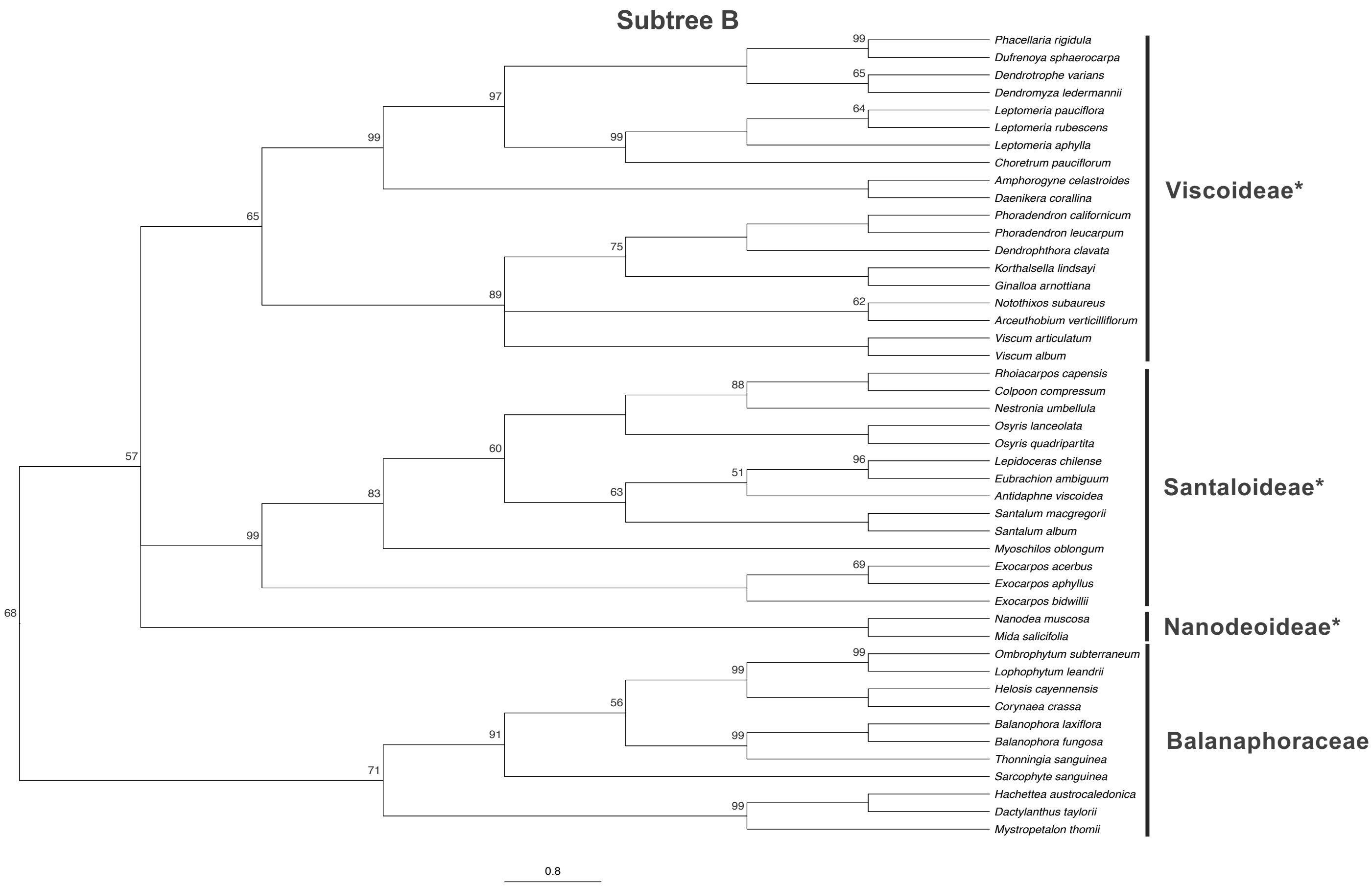

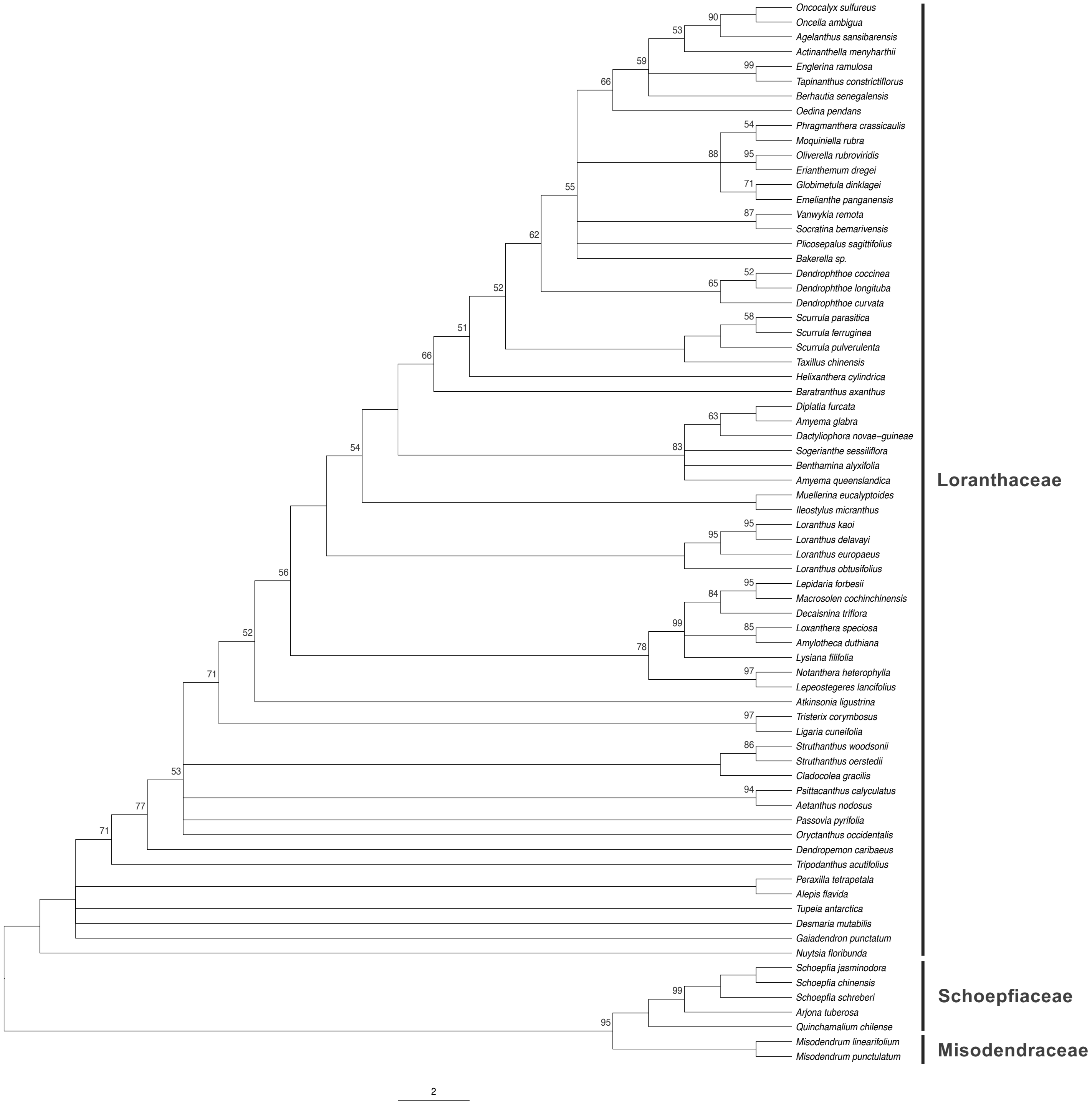

### Fig. S2

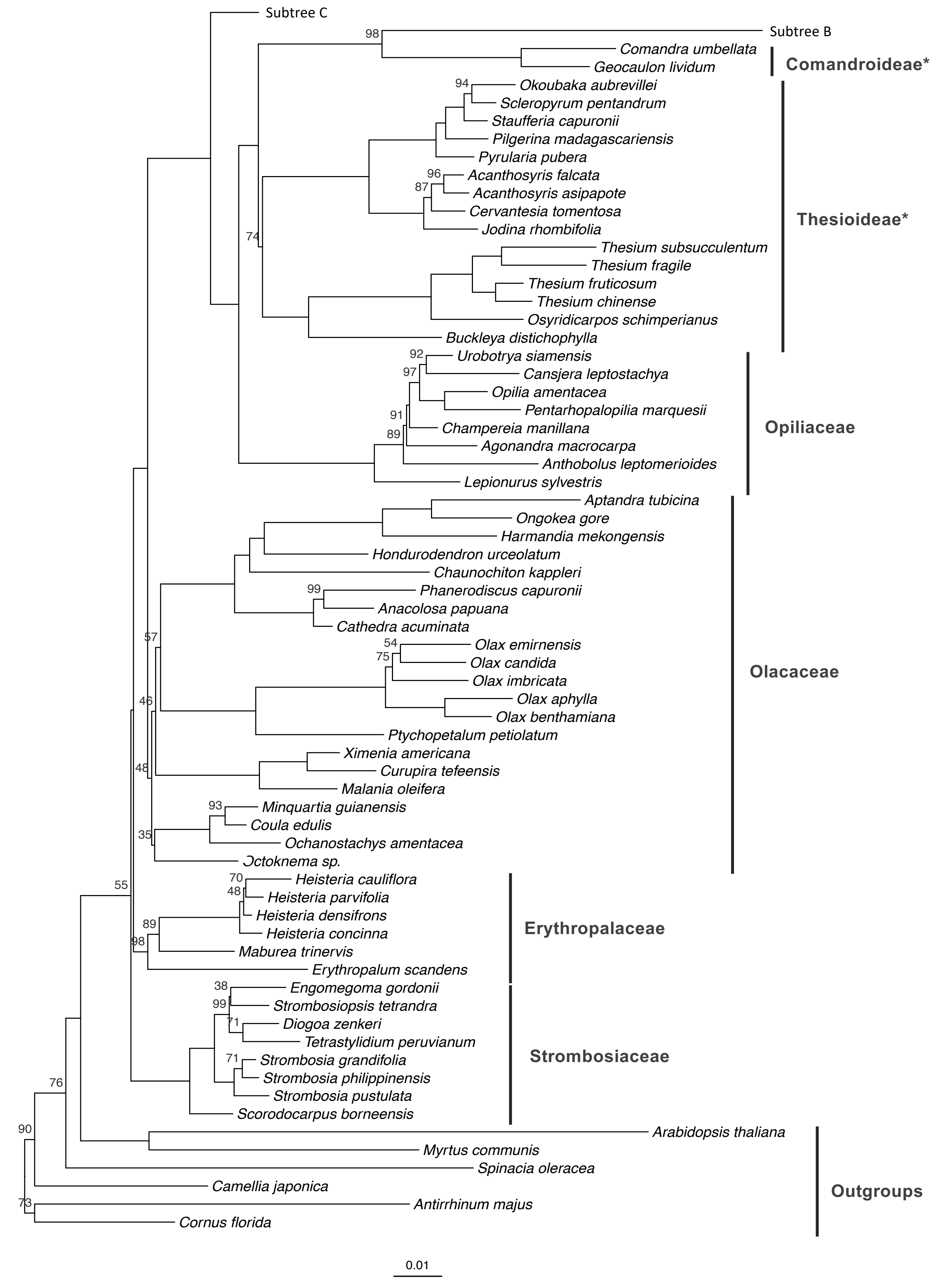

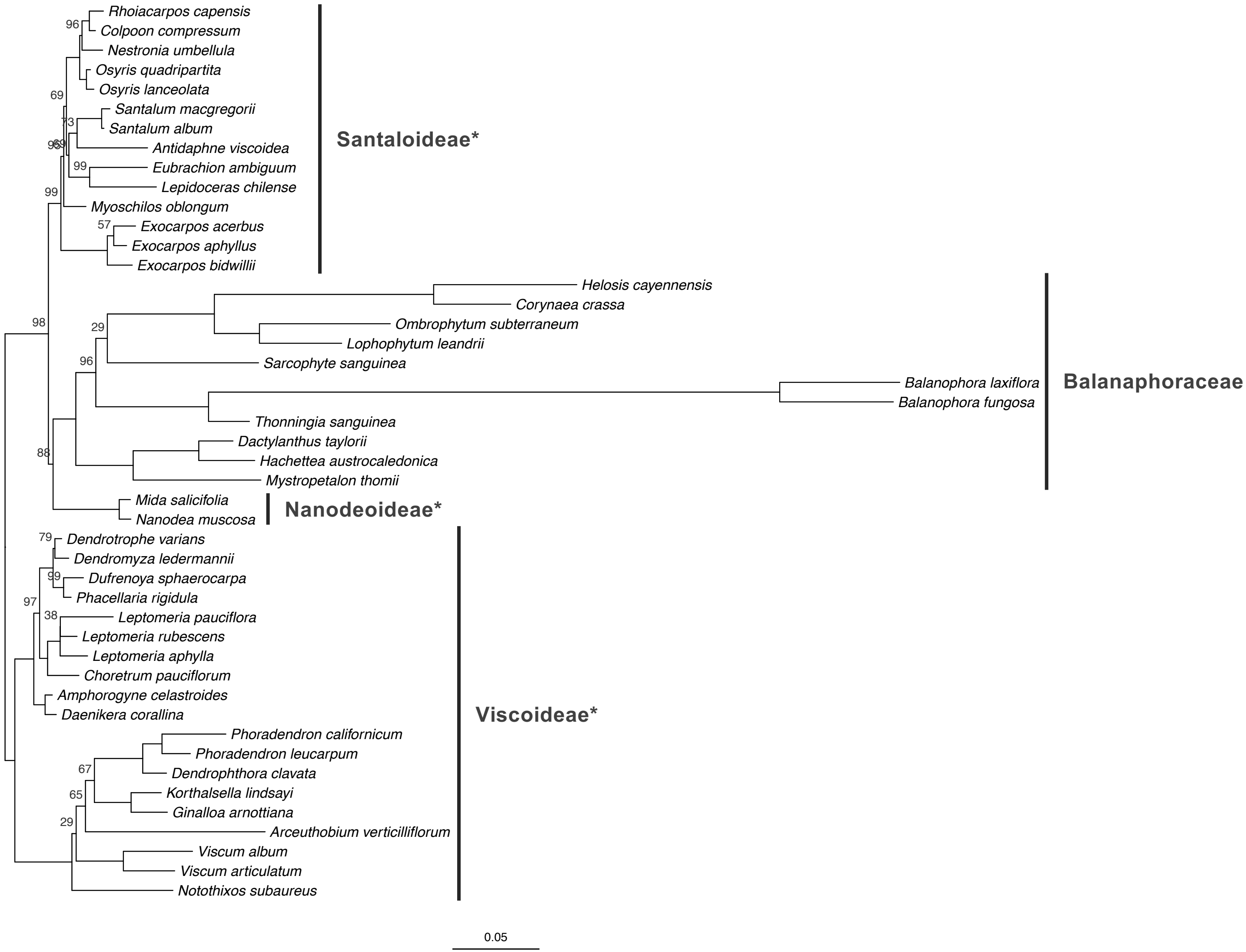

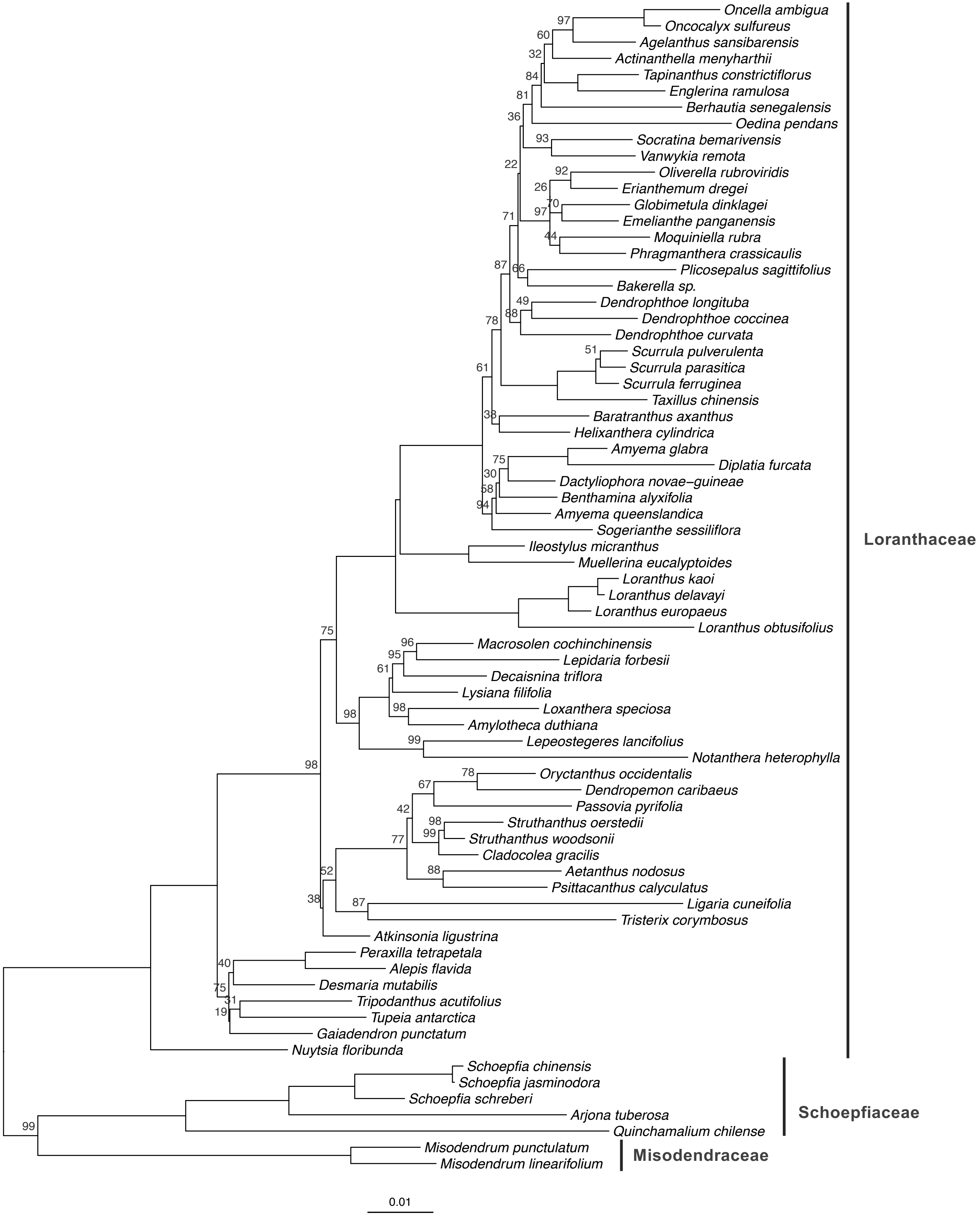
